# scMomer: A modality-aware pretraining framework for single-cell multi-omics modeling under missing modality conditions

**DOI:** 10.1101/2025.08.04.668374

**Authors:** Yuhang Liu, Quan Zou, Ran Su, Leyi Wei

## Abstract

Foundation models offer new opportunities to capture cellular behavior from large-scale single-cell data. However, their development has been greatly constrained due to the limited availability of multi-omics profiles. Consequently, most models are designed for a single modality (e.g. scRNA-seq, or scATAC-seq, etc.), restricting their ability to capture the diversity of heterogeneous biological systems. Here, we introduce scMomer, a modality-aware pretraining framework designed for multi-modal representation learning under missing modality conditions. scMomer adopts a three-stage pretraining strategy that learns unimodal cell representations, models joint representations from multi-omics data, and distills multi-modal knowledge to enable multi-omics-like representations from unimodal input. Its modality-specific architecture and three-stage pretraining strategy enable effective learning under missing modality conditions and help capture cellular heterogeneity. Through extensive experiments, scMomer generates biologically meaningful embeddings and outperforms state-of-the-art unimodal approaches across diverse gene-level and cell-level downstream tasks, including cross-modality translation, gene function prediction, cell annotation, drug response prediction, and perturbation prediction. Overall, these results demonstrate that scMomer serves as a robust, generalizable, and scalable foundation for single-cell multi-modal analysis under missing modality conditions.

## Introduction

Single-cell sequencing technologies have enabled the systematic characterization of complex biological systems at cellular resolution, offering powerful tools to study gene regulation, cellular states, and developmental dynamics^1,2^. Among them, single-cell RNA sequencing (scRNA-seq) captures gene expression patterns^3^, while single-cell ATAC sequencing (scATAC-seq) reveals chromatin accessibility landscapes^4,5^. Recent advances have also made it possible to profile multiple molecular modalities in the same cell, such as gene expression and chromatin accessibility, providing complementary views of cellular regulation^6–8^. However, these multi-omics profiles remain limited in scale and accessibility due to technical and cost constraints, resulting in widespread missing modalities in practice.

Meanwhile, the emergence of foundation models has transformed various domains in artificial intelligence^9–11^. These models leverage large-scale unlabeled data through pretraining to learn general-purpose representations that can be adapted to diverse downstream tasks^12^. In the context of biology, cells can be regarded as basic units of information, with high-dimensional molecular profiles analogous to structured signals in language or vision^13^. Inspired by this analogy, recent studies have begun to explore the use of large pretrained models for single-cell data, aiming to learn transferable representations that improve the accuracy, robustness, and scalability of downstream analyses^14–17^. These approaches typically rely on unsupervised pretraining on large collections of unimodal single-cell data, followed by supervised fine-tuning on specific tasks such as cell type classification, perturbation inference, and drug response prediction.

Despite these advances, existing single-cell foundation models are predominantly limited to a single modality, which restricts their ability to capture the full regulatory landscape of cellular systems^18^. Each omics modality provides a distinct but incomplete view of cellular function; for example, transcriptome profiles reflect gene expression outputs, while chromatin accessibility relates to upstream regulatory potential. Modeling from a single modality may thus miss cross-modality dependencies that are essential for accurately characterizing cellular heterogeneity^19^. A key challenge is to develop models that can learn comprehensive and robust multi-omics cell representations, even when only a single modality is available.

While many recent methods have explored multi-modal representation learning in single-cell analysis, most require all modalities to be present during both training and inference. These mainly include joint embedding frameworks that align multiple omics profiles into a shared latent space^20–23^. For instance, dual-encoder architectures such as scButterfly^21^ project RNA and ATAC data into aligned spaces and reconstruct signals across modalities. Other approaches like scCLIP^22^ and scPair^23^ adopt contrastive learning or feature mapping to learn joint representations. However, these models often assume complete multi-omics input and do not generalize well when some modalities are missing. In addition, many rely on hand-crafted features or modality-specific preprocessing steps, limiting their scalability and generality.

To address missing modality problems, generative methods have been proposed to impute or reconstruct the missing data^24–28^. For example, models such as UnitedNet^25^, MultiVI^26^, scPairing^27^, and scDiffusion-X^28^ learn mappings between modalities using encoder-decoder frameworks or generative diffusion processes. However, most of these efforts focus on data reconstruction rather than learning informative, transferable representations that remain effective under modality dropout. Moreover, they are not designed as foundation models that can generalize across a wide range of tasks and datasets.

Here, we present scMomer, a modality-aware pretraining framework for single-cell multi-omics modeling under missing modality conditions. Unlike previous approaches, scMomer is explicitly designed to learn robust, multi-omics-like representations using unimodal inputs. It adopts a three-stage pretraining strategy: (i) learning unimodal representations via modality-specific encoders with masked modeling objectives; (ii) training on multi-omics data to capture cross-modality interactions; and (iii) distilling knowledge from missing modalities into a compact student encoder that preserves the benefits of multi-omics integration during downstream analysis, even when only a single modality is provided. This design enables scMomer to approximate the performance of full multi-modal models using only unimodal input, addressing a critical limitation of current single-cell foundation models. We evaluate scMomer across multiple single-cell analysis tasks, including multi-sample integration, cross-modality translation, gene function prediction, cell annotation, drug response prediction, and perturbation prediction. The results show that scMomer consistently outperforms state-of-the-art models that rely on unimodal input. These findings demonstrate the potential of scMomer as a general-purpose foundation model for single-cell analysis.

## Results

### The scMomer pretraining framework

scMomer is a modality-aware pretraining framework designed to enable robust representation learning for single-cell multi-omics data under missing modality conditions (Fig. 1a). While retaining the strengths of existing foundation models, scMomer introduces a structured three-stage training strategy to learn unimodal, cross-modal, and modality-agnostic representations through knowledge distillation. This training strategy allows scMomer to generalize effectively when a single modality is available, a common limitation in single-cell analysis.

**Figure 1.**
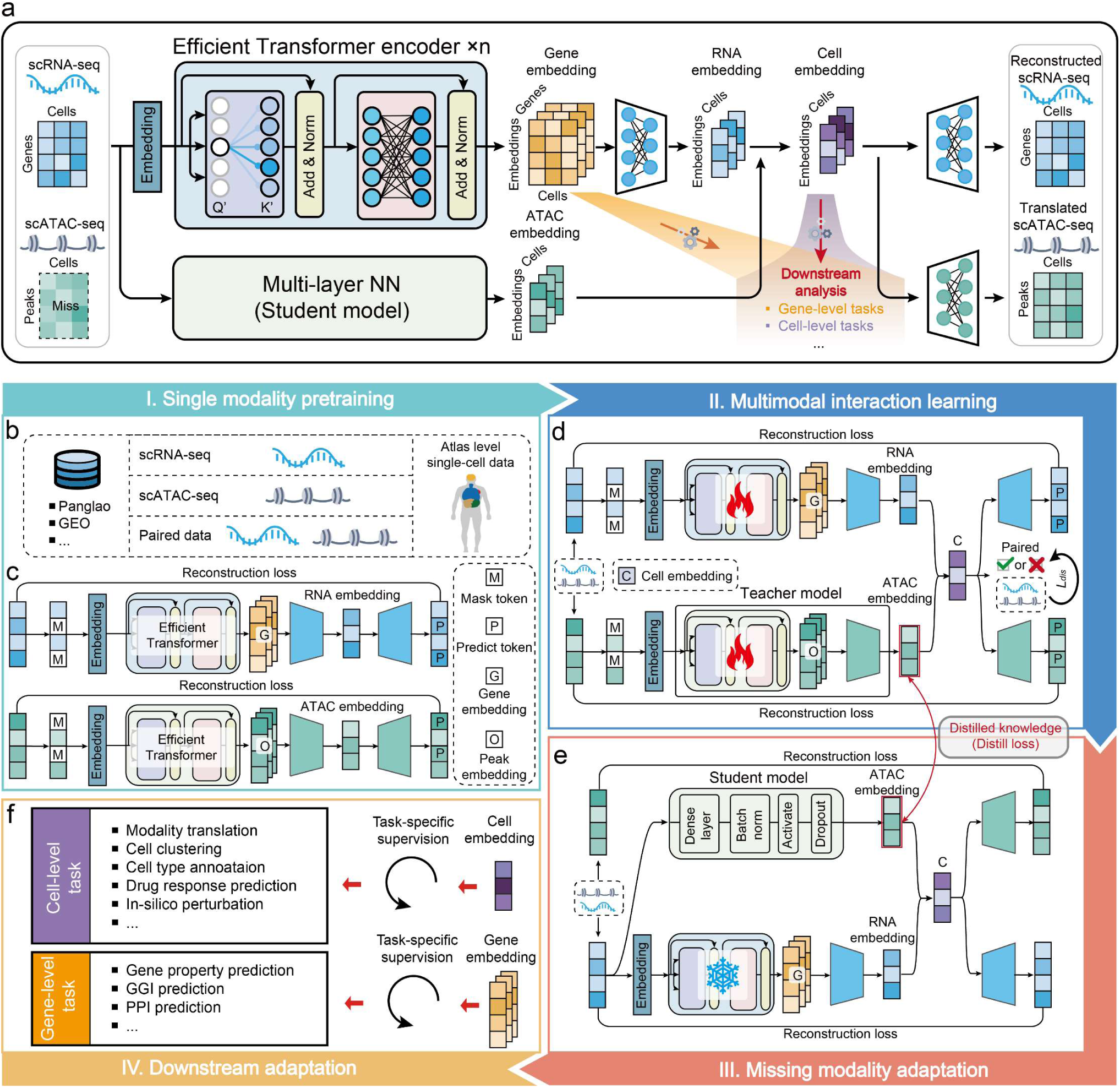
Overview of the scMomer pretraining framework for single-cell multimodal analysis with missing modality. **a,** Workflow of scMomer in the presence of missing modality. The model takes single-modality data (e.g., scRNA-seq) as input and generates comprehensive cell embeddings. The architecture consists of efficient Transformer blocks for learning RNA-based representations and a multi-layer neural network that infers ATAC-based embeddings through knowledge distillation. The RNA and ATAC embeddings are integrated to form a unified cell embedding, which can be used to reconstruct missing modalities (e.g., scATAC-seq) and enable downstream analyses. **b,** Examples of datasets used in the pretraining stage, including scRNA-seq, scATAC-seq, and multimodal data from atlas-level resources such as Panglao and GEO. **c,** Stage I: Single-modality pretraining. The model is trained on large-scale independent scRNA-seq and scATAC-seq data using masked token prediction and reconstruction loss to learn gene- and peak-level embeddings. **d,** Stage II: Multimodal interaction learning. Pretrained model is fine-tuned on multimodal dataset to learn the relationships between RNA and ATAC modalities using reconstruction loss and a modality-pairing discrimination loss. The learned ATAC embeddings are used as distilled knowledge. **e,** Stage III: Missing modality adaptation. A lightweight student model is trained to reconstruct ATAC embeddings from RNA inputs using distillation loss, enabling robust embedding learning when one modality is missing. **f,** The pretrained representations can be adapted to a range of cell-level (e.g., modality translation, cell annotation, drug response prediction, and perturbation analysis) and gene-level tasks (e.g., gene property prediction, gene–gene interaction inference, and protein–protein interaction prediction).

In the first stage, we adopt masked modeling strategies to capture intra-modality interactions (Fig. 1b,c). For RNA, this involves predicting the expression of masked genes based on the context of co-expressed genes within the same cell. For ATAC, the model predicts the accessibility of masked chromatin patches using information from surrounding genomic regions. This strategy allows the model to learn intrinsic gene–gene interactions and chromatin accessibility patterns from unimodal input. In the second stage, the pretrained unimodal encoders are jointly fine-tuned using pseudo-paired RNA–ATAC profiles collected from atlas-scale datasets (Fig. 1d). RNA and ATAC embeddings are fused to learn unified cell representations that capture complementary signals across modalities. To improve cross-modal alignment, we introduce a modality-matching discriminator, which encourages the model to distinguish between correctly and incorrectly paired inputs^29^. This facilitates more coherent and biologically relevant integration of multi-omics data. In the final stage, we introduce a compact student model based on feedforward networks (FFNs), motivated by prior studies like scPair^23^ that demonstrate their effectiveness in cross-modal translation. The student is trained to approximate the missing modal embeddings produced by the modality-specific encoder, enabling inference when only one modality is available (Fig. 1e). Through this distillation process, scMomer effectively learns modality-agnostic representations that retain meaningful biological information, even under missing modality conditions. After training, the representations learned by scMomer can be applied to a wide range of downstream tasks in single-cell analysis, including cross-modality translation, cell annotation, drug response prediction, perturbation modeling, and so on (Fig. 1f).

To maximize compatibility and efficiency, we build scMomer upon existing foundation models rather than designing new architectures from scratch. This approach offers two main advantages. First, it leverages the strengths of well-established models for individual modalities, reducing computational overhead. Second, the modular architecture of scMomer allows for flexible replacement of each encoder as more advanced models and larger datasets become available, making it a general and extensible framework for multi-omics pretraining. Specifically, we use pretrained scBERT^30^ as the RNA encoder and adopt a ViT-based ATAC encoder^31^, which has been shown effective in scCLIP^22^. This design ensures both practical usability and future extensibility across different data scenarios.

### scMomer supports multi-omics integration and cross-modality translation

Unlike conventional single-cell multi-omics integration methods that primarily aim to align different modalities from the same cell into similar representations, we propose an alternative strategy. scMomer learns modality-specific cell embeddings independently and then fuses them into a unified, comprehensive cell representation. To assess this approach, we first evaluated the representation quality of scMomer embeddings derived from single modalities (RNA or ATAC), as well as the integrated multi-modal embeddings. We further investigated the model’s capacity for cross-modality translation.

To evaluate the zero-shot representation capacity and integration ability of scMomer, we applied the pretrained model to the human brain dataset from 10x Genomics, generating RNA embeddings, ATAC embeddings, and integrated cell embeddings. We aimed to assess whether scMomer can learn modality-specific cell representations and synthesize them into robust and biologically informative unified embeddings, even in missing modality scenarios. As shown in Fig. 2a, UMAP visualization of the learned embeddings revealed clearer biological structure compared to the raw input data. This was especially evident for ATAC data, where raw features exhibited weak cell-type separation, while scMomer’s inferred ATAC embeddings maintained distinct clusters. Although the RNA and ATAC embeddings exhibited distinct spatial distributions (Fig.2b), both preserved modality-specific biological information. This suggests that scMomer captures complementary modality-specific features rather than enforcing alignment.

**Figure 2.**
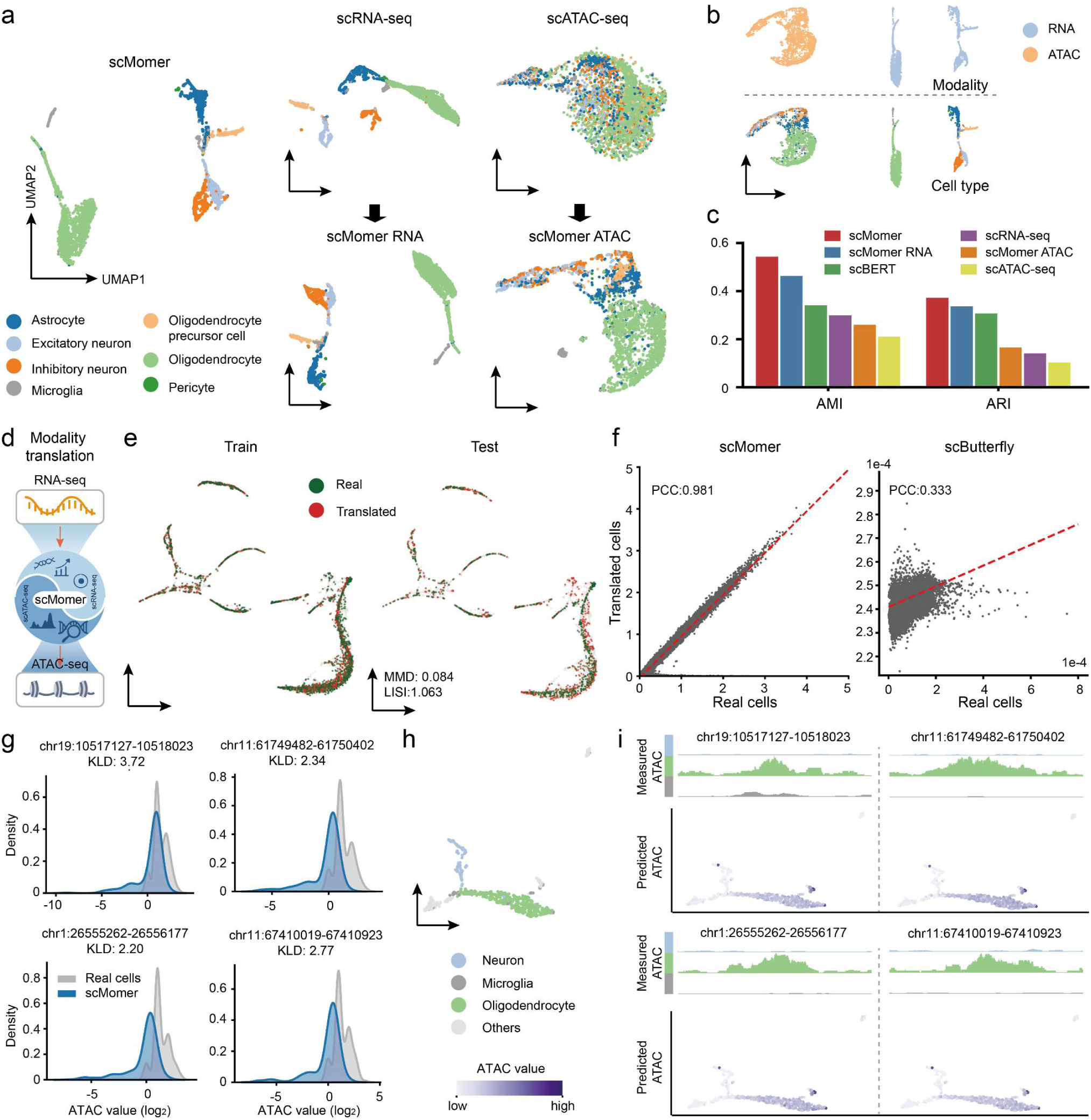
scMomer enables integrative multi-omics representation and cross-modality translation. **a,** UMAP visualization for different embeddings on the human brain dataset. **b,** UMAP visualization for RNA and ATAC joint embeddings. Cells are colored by modality (upper) and by cell type (lower). **c,** Comparison of embedding performance across six methods. **d,** Schematic overview of cross-modality translation using scMomer, where RNA input is used to infer missing ATAC modality. **e,** UMAP visualization comparing scMomer-translated and ground truth scRNA-seq data in both training and test sets. **f,** Correlation plot comparing averaged translated ATAC and observed ATAC accessibility across peaks, simulating bulk-level signals. Each dot represents a chromatin peak. scButterfly operates on normalized data. **g,** Distribution of the true and predicted chromatin accessibility profiles for the top four differential peaks in the Oligodendrocyte cell type. **h,** UMAP of cells on the test set, colored by cell groups. **i,** Comparison of true chromatin accessibility levels across cell types for differentially accessible regions (upper). UMAP visualization of the translated accessibility profiles corresponding to differential peaks (lower), with color indicating predicted chromatin opening probability.

To further validate this observation, we performed k-means clustering (*k* = 7, based on cell type count) on the raw features and the learned embeddings. Both RNA and ATAC embeddings generated by scMomer achieved higher Adjusted Mutual Information (AMI) and Adjusted Rand Index (ARI) scores than the original data, and the fused cell embeddings achieved the highest performance (Fig. 2c), confirming the benefit of modality-aware representation learning.

Next, we evaluated scMomer’s ability to perform cross-modality translation (Fig. 2d), a challenging task in single-cell multi-omics analysis. We fine-tuned scMomer on the human brain dataset and compared it with scButterfly^21^, a recent method designed for cross-modality translation. As shown in Fig. 2e, UMAP visualizations revealed that the scMomer-translated scATAC-seq data closely resembled real ATAC profiles. Additionally, we computed pseudo-bulk profiles by averaging ATAC peaks across translated cells and compared them with real bulk profiles. As shown in Fig. 2f, scMomer achieved a Pearson Correlation Coefficient (PCC) of 0.981, significantly surpassing scButterfly’s 0.333. The need for TF-IDF normalization in scButterfly may constraint its ability to recover true accessibility values, limiting population-level consistency.

We further examined how well the translated ATAC profiles recovered differential peaks. Fig. 2g shows that the distribution of marker peaks in scMomer-translated ATAC data more closely matched that of the real ATAC data. These results suggest that, unlike traditional RNA-only foundation models that primarily learn gene–gene interactions, scMomer captures inter-modality relationships and enables effective cross-modality translation. At the single-locus level, we highlighted the differential peaks in Fig. 2g, whose accessibility patterns were identified as cell type markers (Fig. 2i)^32^. These peaks exhibited predicted accessibility patterns in the held-out samples (Fig. 2i) that were consistent with their corresponding cell types (Fig. 2h).

In summary, these findings demonstrate that scMomer can learn meaningful interactions across multi-modalities and effectively generalize to cross-modality integration and translation tasks, supporting its potential in broader multi-omics applications.

### scMomer enables multi-sample integration while preserving cell biology

In this section, we demonstrate that the zero-shot scMomer embedding can integrate multi-batches based on the principles of cell biology, resulting in cells with similar cell types or disease states having more similar representations (Fig. 3a). We evaluated scMomer embeddings on two single-cell RNA datasets representing immune cells from the aorta and cardiomyocyte, including both healthy and diseased samples.

**Figure 3.**
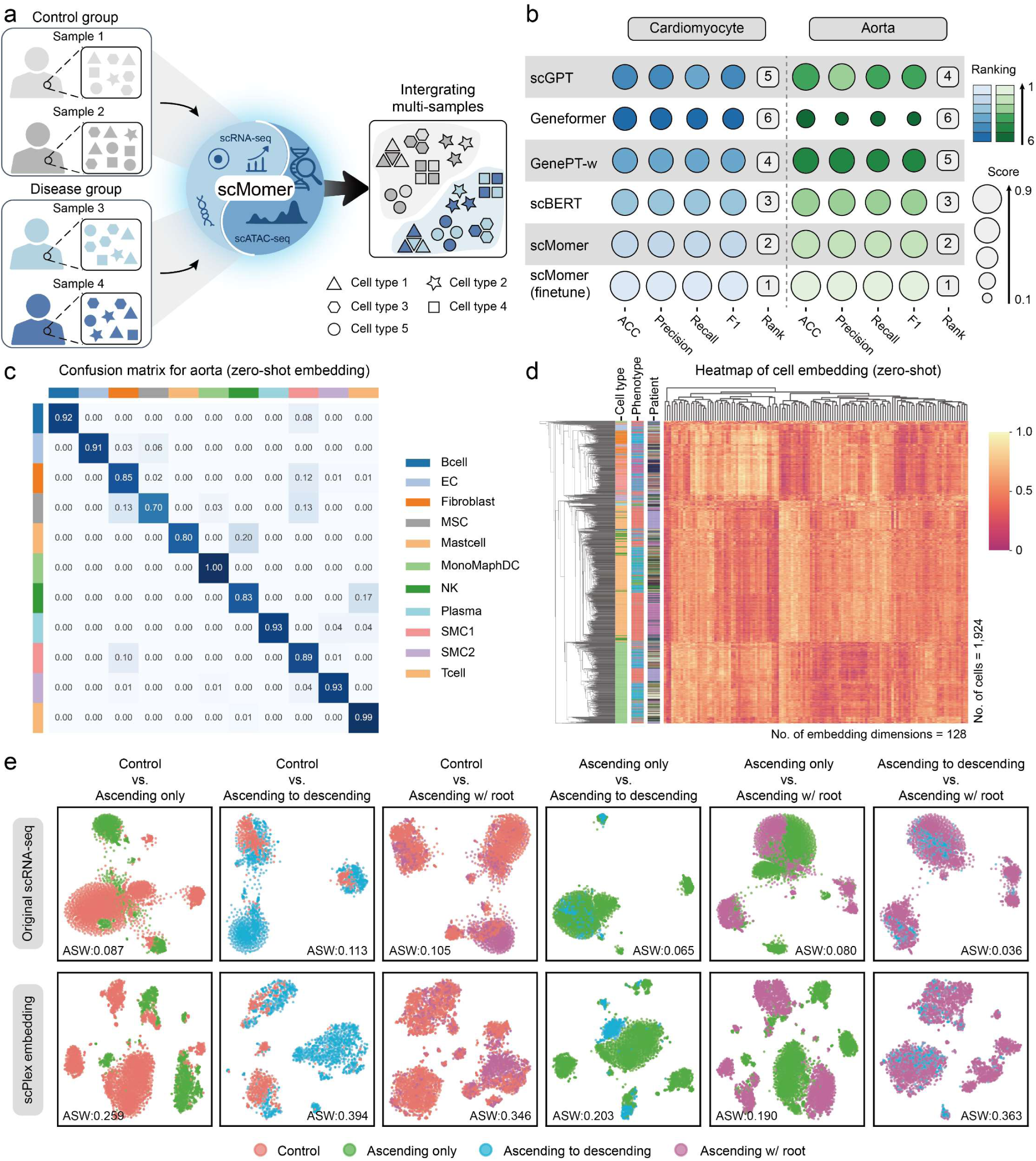
scMomer integrates multi-batch while preserving biological information. **a,** Illustration of the scMomer for multi-batch integration. **b,** The performance of different methods on disease/phenotype prediction tasks. The blue and green bubbles represent the performance on cardiomyocyte and aorta datasets, respectively. The lighter the color, the better the performance. **c,** Confusion matric of the zero-shot cell embedding in cell type prediction on the aorta dataset. The confusion matrices of other methods are included in Supplementary Fig. 5. **d,** Hierarchical clustering of zero-shot scMomer aorta cell embeddings. **e,** t-SNE of the aorta dataset, coloured by four different phenotypes. The visualization plots of other methods are included in Supplementary Fig. 1.

We first evaluated the discriminative capacity of zero-shot cell embeddings in capturing biologically relevant annotations, including cell type and disease phenotype. Specifically, we compared embeddings generated from five pretrained models: (i) scGPT, (ii) Geneformer, (iii) GenePT-w, (iv) scBERT, and (v) scMomer. We evaluated performance by training L2-regularized logistic regression classifiers on 80% of the data and testing on the remaining 20%, using the embeddings as input features. scMomer achieved higher accuracy, precision, recall, and F1 scores compared to baseline models on the aorta and cardiomyocyte datasets for phenotype classification (Fig. 3b, and Supplementary Fig. 3,4).

We further investigated whether fine-tuning with limited task-specific data could improve accuracy. After fine-tuning, scMomer achieved a notable increase in performance, reaching approximately 0.95 accuracy on both datasets (Fig. 3b). For the aorta dataset, improvements were mainly observed in distinguishing between the “ascending to descending” and “ascending with root” subtypes, which may involve inflammatory responses and gene expression changes that complicate differentiation. In the cardiomyocyte dataset, gains were largely attributed to better separation of non-failing heart (NF) and hypertrophic cardiomyopathy (HCM), which aligns with clinical observations that genetic mutations can drive progression from asymptomatic to HCM states^33^.

Using the same evaluation strategy, we assessed cell type classification based on zero-shot embeddings. Results in Fig. 3c and Supplementary Fig. 5 demonstrate that pretrained embeddings from scBERT, scGPT, GenePT-w, and scMomer all achieved reasonable performance. Notably, scMomer outperformed other models, particularly in identifying rare cell types with fewer training samples. For example, in the aorta dataset, baseline methods struggled with mast cells, B cells, and endothelial cells, which are the least represented (Supplementary Fig. 5). In contrast, logistic regression using scMomer embeddings achieved accuracies of 0.80, 0.92, and 0.91 for these cell types.

We then evaluated the biological coherence of the resulting clusters in terms of both cell type and disease phenotype. Fig. 3e shows t-SNE visualizations of the raw gene expression data (top) and the scMomer zero-shot embeddings (bottom), both colored by patient phenotype. Compared to raw data, the scMomer embeddings more clearly separated distinct phenotypes. For instance, “ascending only” and “ascending to descending” cases were well-separated by scMomer (green and blue points, respectively). In addition, we visualized clustering results on both cell type and patient identity using k-means and quantified their alignment with true labels. Similar to phenotype classification, scMomer embeddings better distinguished cell types and aligned patient-level batch effects. For the aorta dataset, the ARI for k-means clustering on scRNA-seq data was 0.1214 (cell type) and 0.0734 (phenotype), while scMomer achieved 0.1319 and 0.3699, respectively (Supplementary Fig. 1). For comparison, scBERT embeddings resulted in ARIs of 0.0659 and 0.1628. For clustering by patient identity, raw scRNA-seq data showed an ARI of 0.3014, whereas scMomer achieved a slightly lower ARI of 0.2271, indicating reduced sensitivity to batch effects. Similarly, scMomer also outperformed scBERT on the cardiomyocyte dataset (Supplementary Fig. 2). These findings suggest that scMomer embeddings preserve meaningful biological signals and exhibit greater robustness to batch effects.

### scMomer learns gene embeddings for gene-level tasks

We next evaluated whether gene embeddings derived from scMomer can support various gene-level downstream applications. For each input scRNA-seq profile, scMomer generates a 200-dimensional embedding for every gene, capturing gene-specific features in a given cellular context. To obtain a unified gene representation, we averaged gene embedding across all cells from the pretrained model. Using these averaged gene embeddings, we assessed performance on several benchmark tasks, including gene property prediction, and gene–gene/protein–protein interaction prediction.

We first tested scMomer on four gene property classification tasks: (i) dosage-sensitive vs. insensitive transcription factors (TFs), (ii) bivalent vs. non-methylated genes, (iii) bivalent vs. Lys4-methylated genes, and (iv) long-range vs. short-range TFs. As baselines, we included GenePT^34^, scBERT^30^, Gene2vec^35^, and randomly embeddings as a control. To ensure fair comparison despite differences in gene vocabularies across models, we used the intersection of covered genes from the original study^15^. All methods were evaluated via five-fold cross-validation. We directly used the pretrained gene embeddings (Supplementary Fig. 6) with two standard classifiers: L2-regularized logistic regression and random forest. As shown in Table 1, scMomer consistently outperformed the unimodal baseline scBERT, achieving AUC improvements of 0.018, 0.036, 0.004, and 0.004 across the four tasks. Performance was also competitive with or superior to other embedding-based models. These improvements likely stem from scMomer’s ability to capture interactions across modalities during pretraining, allowing more accurate gene-level representations. Notably, GenePT outperformed other methods on tasks (i) and (ii), suggesting that integrating prior knowledge of gene function from text may offer a promising direction for further improvement.

**Table 1.** Five-fold cross validation for scMomer along with other approaches on the four binary gene property classification tasks.

| Method | Fivefold cross validation AUC ± s.d. | | | |
| --- | --- | --- | --- | --- |
|  | Long vs. short TF range | Dosage sensitivity | Bivalent vs. non-methylated | Bivalent vs. Lys4-methylated |
| Random embed + LR | 0.553 ± 0.067 | 0.507 ± 0.055 | 0.485 ± 0.112 | 0.502 ± 0.016 |
| Random embed + RF | 0.584 ± 0.089 | 0.501 ± 0.083 | 0.529 ± 0.116 | 0.490 ± 0.006 |
| Gene2vec + LR | 0.613 ± 0.142 | 0.815 ± 0.037 | 0.644 ± 0.027 | 0.763 ± 0.073 |
| Gene2vec + RF | 0.549 ± 0.132 | 0.838 ± 0.120 | 0.556 ± 0.062 | 0.790 ± 0.014 |
| GenePT+LR | 0.614 ± 0.096 | 0.798 ± 0.128 | 0.468 ± 0.155 | 0.675 ± 0.042 |
| GenePT+RF | **0.657 ± 0.074** | **0.870 ± 0.139** | 0.473 ± 0.101 | 0.645 ± 0.040 |
| scBERT + LR | 0.621 ± 0.142 | 0.812 ± 0.072 | 0.657 ± 0.115 | 0.776 ± 0.063 |
| scBERT + RF | 0.561 ± 0.137 | 0.822 ± 0.101 | 0.553 ± 0.098 | 0.799 ± 0.013 |
| scMomer + LR | 0.639 ± 0.129 | 0.857 ± 0.028 | **0.661 ± 0.052** | 0.763 ± 0.069 |
| scMomer + RF | 0.573 ± 0.131 | 0.858 ± 0.083 | 0.621 ± 0.109 | **0.803 ± 0.013** |

\* For the Bivalent vs. Non-methylated prediction task, we used all bivalent regions from a single chromosome and genome-wide non-methylated regions to simulate an imbalanced dataset. For the Bivalent vs. Lys4-methylated task, genome-wide data were used for training.

We further evaluated gene embedding performance in gene–gene interaction (GGI) prediction. Following Du et al.^35^, we used GO-based annotations to define gene pairs, concatenated their embeddings, and applied logistic regression for classification. As shown in Supplementary Table 1, scMomer achieved an AUC of 0.787, outperforming scBERT by 0.766 and exceeding other methods.

In addition, we assessed scMomer’s ability to predict protein–protein interactions (PPI). Following the procedure of Chen et al.^34^, we mapped protein IDs to gene names using UniProt^36^, concatenated the embeddings of gene pairs, and used logistic regression for prediction. Evaluation on the HuRI dataset^37^ showed that scMomer achieved an AUC of 0.636, outperforming the scBERT baseline by 0.632 (Supplementary Table 1). Overall, these results highlight the effectiveness of scMomer in gene-level tasks, attributed to its modality-aware pretraining that captures modality cross-talk beyond unimodal approachs.

Beyond global gene embeddings, we also explored cell-context-specific variation in gene representations. We quantified this by measuring the standard error of gene embeddings across cell types from aortic tissue in control individuals. As expected, housekeeping genes such as *GAPDH* showed low variability across cell types, while context-dependent genes like *NOTCH* receptors exhibited higher variability (Supplementary Fig. 7), further supporting the biological validity of scMomer’s learned representations.

### scMomer embeddings enhance cell type annotations

We next examined scMomer’s utility in cell type annotation, which is one of the central tasks in single-cell analysis. We performed comprehensive evaluations on five datasets: Zheng68K^38^, Baron^39^, Segerstolpe^40^, Muraro^41^, and Xin^42^. Zheng68K is a widely used PBMC dataset. The remaining four are pancreas datasets, representing diverse human cell types and sequencing technologies. This makes them ideal for evaluating model generalizability. Following prior studies on foundation models, we trained a multi-layer perceptron (MLP) classifier using the pretrained embeddings. Most foundation model parameters were frozen during finetuning. We considered two evaluation scenarios: intra-dataset and out-of-distribution validation. Baseline methods included CellID^43^, sciBet^44^, SingleR^45^ and scBERT, with scBERT served as our primary unimodal baseline.

#### Intra-dataset validation

For in-distribution evaluation, we applied five-fold cross-validation to split each dataset into training, validation, and test sets. As shown in Fig. 4a, scMomer surpassed the competing methods in both accuracy and macro F1-score on most of the datasets. On the challenging Zheng68K dataset, scMomer reached an accuracy of 0.808 and F1-score of 0.703, outperforming the scBERT (accuracy = 0.768, F1 = 0.647). Notably, scMomer significantly improved classification of hard-to-distinguish cell types such as CD8+ cytotoxic T cells and CD8+/CD45RA+ T cells, with respective gains of 0.820 and 0.870 in accuracy (Fig. 4b,c, and Supplementary Figure 8). These results indicate that scMomer’s multi-modal pretraining enhances cell representation quality, particularly for closely related cell types^39^.

**Figure 4.**
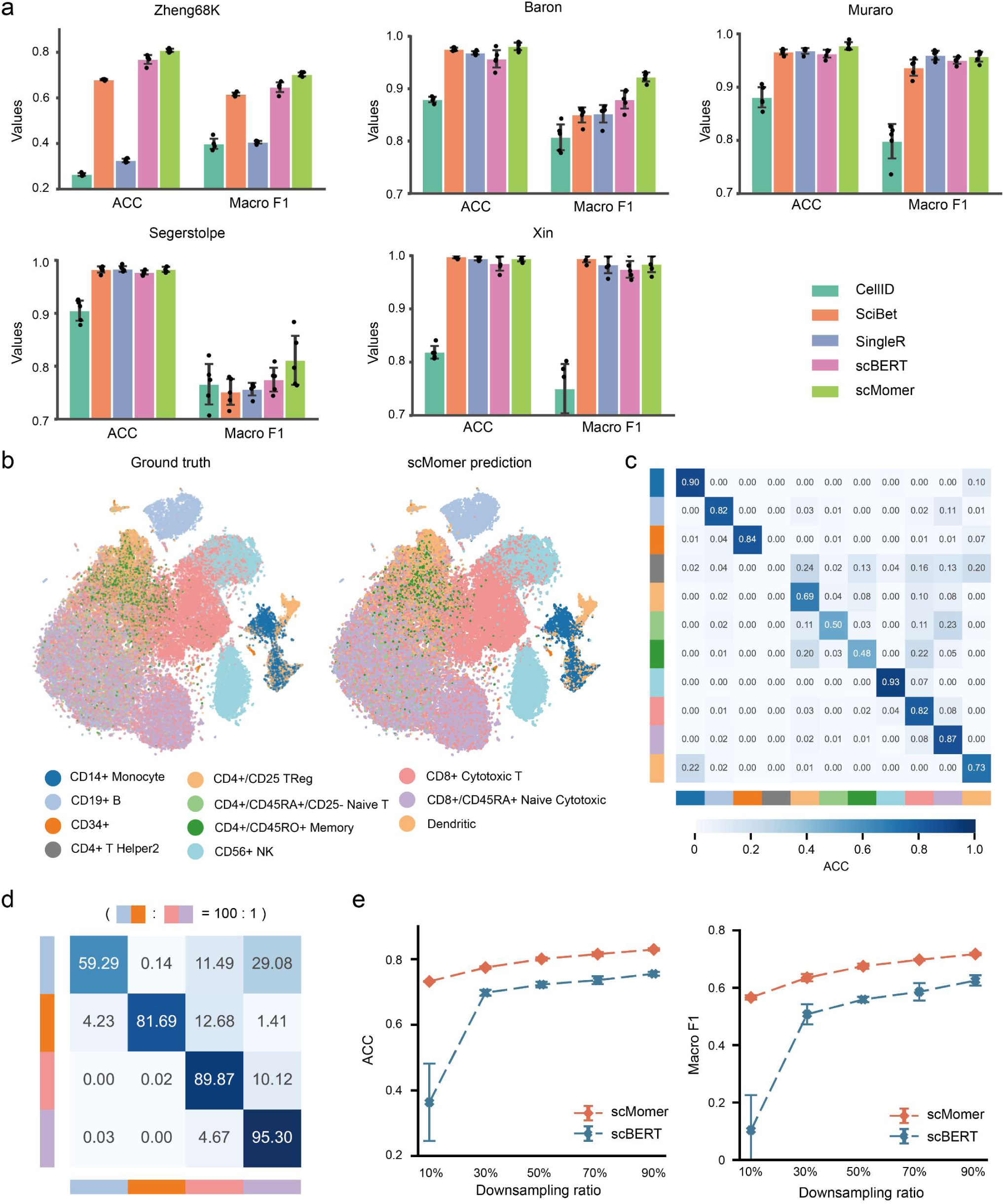
Universality of scMomer across diverse datasets. **a,** Cell type annotation accuracy on five benchmark datasets using fivefold cross-validation. Box plots indicate the median, interquartile range, and 1.5× interquartile range of accuracy across all methods. **b,** t-SNE visualization of the Zheng68K dataset. Left: true cell type labels; right: predicted labels by scMomer. Predictions are aggregated across test folds from fivefold cross-validation (n = 68,450 cells). The t-SNE plots of competing methods are shown in Supplementary Fig. 9. **c,** Confusion matrix of scMomer predictions on the Zheng68K dataset using fivefold cross-validation. **d,** Performance of scMomer under imbalanced conditions, evaluated at a cell type imbalance ratio of 100:1. **e,** Model performance across varying dataset sizes. The x-axis denotes the downsampling proportion from the original Zheng68K dataset, and y-axis shows cell type annotation accuracy.

We also evaluated model performance on imbalanced datasets. We constructed a synthetic imbalance in Zheng68K with a 100:1 ratio between abundant and rare cell types. As shown in Fig. 4d, scMomer maintained robust performance with an accuracy of 0.838 and a macro F1 score of 0.829, surpassing scBERT by 0.023 and 0.033, respectively, and significantly surpassing classical annotation tools (Supplementary Table 2). These results suggest that scMomer is robust to data imbalance, likely due to its capacity to learn generalized representations from multi-modal pretraining. To assess scalability, we downsampled Zheng68K dataset to create subsets containing 10% to 90% of the original data. As shown in Fig. 4e, scMomer consistently outperformed scBERT across all subset sizes, with especially large margins when training data was limited (∼10%).

#### Out-of-distribution validation

To evaluate generalization across datasets and technologies, we tested zero-shot transferability. We trained a model on the Baron dataset and tested on the remaining pancreas datasets (Segerstolpe, Muraro, Xin), focusing on four shared cell types (alpha, beta, delta, gamma). Despite substantial batch effects (Supplementary Fig. 8), both scMomer and scBERT showed strong performance (accuracy = 0.962 and 0.955, respectively), outperforming CellID, sciBet, and SingleR (Supplementary Table 3). Overall, these results suggest that its cross-modal knowledge supports more accurate and transferable cell representations.

### scMomer embeddings empowers for drug response prediction

Accurately predicting cancer drug response (CDR) is essential for understanding tumor biology and guiding therapeutic strategies. Two major tasks in this field are drug sensitivity prediction and drug synergy prediction, both of which involve modeling the effects of compounds on cellular systems by integrating molecular structure and multi-omics information. While graph-based methods for molecular representation have matured, generating robust and informative cellular embeddings remains a significant challenge due to the high dimensionality and heterogeneity of omics data. scMomer addresses this challenge by leveraging its modality-aware architecture and pretraining strategy to generate comprehensive cell representations. Although pretrained on single-cell data, we hypothesize that scMomer embeddings can also be transferred to bulk-level drug response prediction, as bulk measurements reflect the “averaged” signal from large numbers of single cells.

To evaluate the utility of scMomer in this setting, we incorporated it into the DeepCDR framework^46^ by replacing the original transcriptomic encoder with scMomer-generated embeddings. Focusing solely on gene expression features for fair comparison, we benchmarked scMomer against DeepCDR and scBERT. As shown in Fig. 5b, scMomer achieved a PCC of 0.907 between predicted and true IC50 values, outperforming DeepCDR and scBERT by margins of 0.053 and 0.060, respectively (Supplementary Fig. 10a). This improvement can be attributed to the reduced overfitting observed in scMomer, likely due to its integration of multi-omics priors during pretraining.

**Figure 5.**
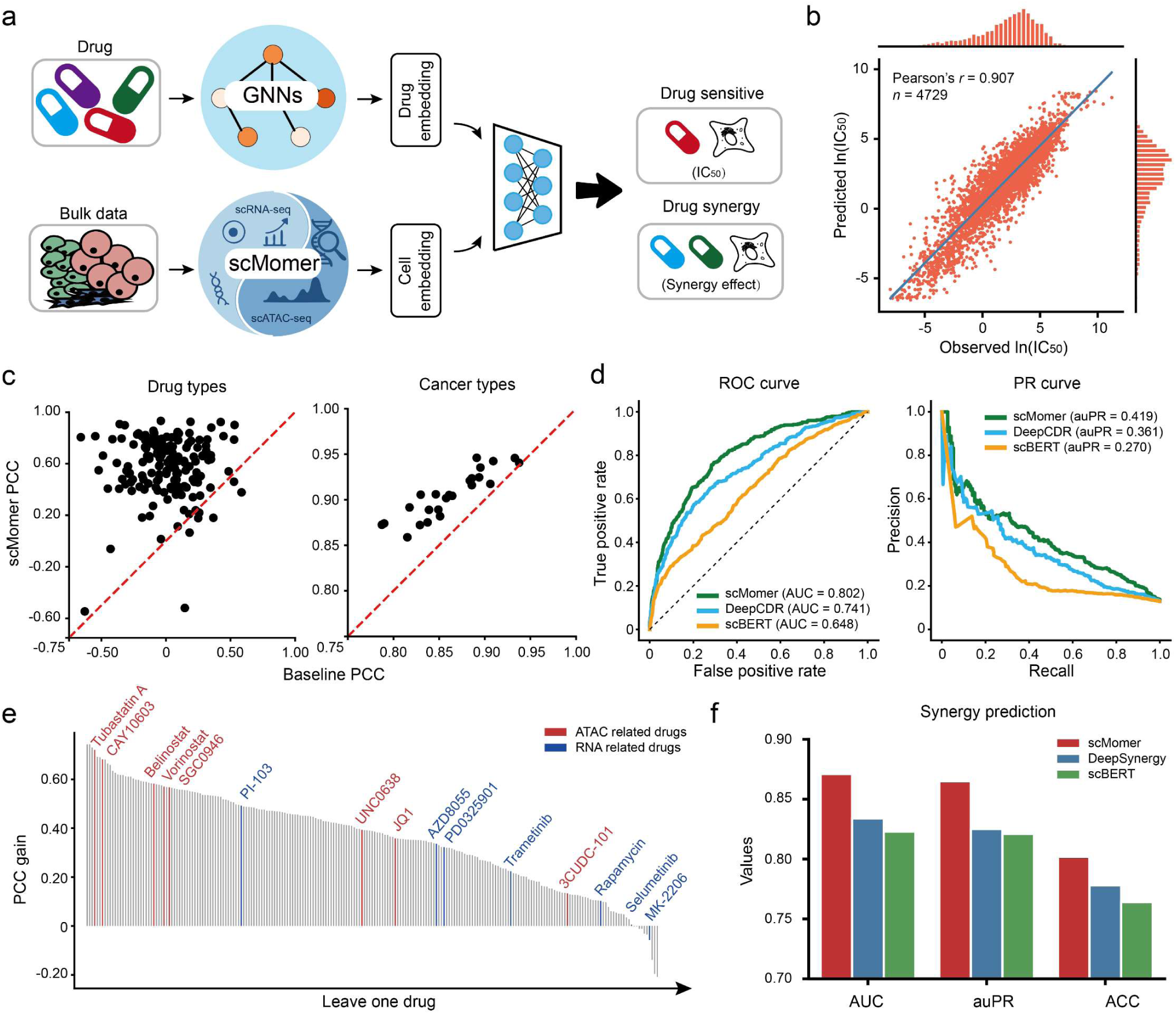
Predicting drug response using scMomer embeddings. **a,** Schematic overview of drug response prediction using scMomer-derived cell representations. **b,** Comparison of predicted and observed IC50 in the test set. **c,** Comparison of PCC for all drug–cancer type pairs in the test set. Each dot represents a drug or cancer type. The x axes show PCCs obtained using the baseline model and the y axes are from scMomer-based model. **d,** AUROC and AUPRC curves for different models in predicting CDR. **e,** Leave-one-drug-out performance of different models. Bars represent PCC gains achieved by scMomer over the baseline for each held-out drug. The red color indicates drugs related to RNA, while the blue color indicates drugs related to ATAC. **f,** Performance comparison across methods for drug synergy prediction.

We further analyzed performance across 30 TCGA cancer types and 223 individual drugs. scMomer consistently outperformed baseline models in the majority of cancer types and drug categories (Fig. 5c and Supplementary Fig. 10b). Notably, the largest performance gain was observed for the MEK inhibitor Trametinib, which blocks the ERK/MAPK pathway through inhibition of MEK1 and MEK2. Given that ERK signaling has been shown to regulate chromatin accessibility^47^, this result further supports the biological relevance of the multi-omics knowledge captured by scMomer.

To evaluate generalization, we conducted a leave-one-drug-out test across all 223 compounds. scMomer outperformed baseline method on 213 drugs, achieving an average PCC improvement of 0.362 (Fig. 5e). We observed that drugs associated with chromatin accessibility, such as histone deacetylase (Vorinostat, Belinostat, and Tubastatin) exhibited greater performance gains compared to RNA-related compounds, such as tyrosine kinase or receptor inhibitors (Fig. 5e). This observation further supports the ability of scMomer to effectively capture and leverage ATAC-related information.

We additionally binarized the IC50 values based on thresholds provided by Iorio et al.^48^, yielding a highly imbalanced dataset (positive to negative ratio ∼7:1). Even under these conditions, scMomer achieved notable gains over competing methods, improving AUC and auPR by 0.061 and 0.154 compared to DeepCDR, and by 0.058 and 0.169 compared to scBERT (Fig. 5d). Finally, we explored the utility of scMomer embeddings for drug synergy prediction. As shown in Fig. 5f, scMomer outperformed both DeepSynergy and scBERT, achieving AUC gains of 0.037 and 0.048, respectively.

Overall, these results highlight the potential of scMomer for drug response modeling and its capacity to enhance understanding of therapeutic mechanisms in cancer. With further integration of advanced methods, scMomer could be extended to support single-cell-level drug discovery^49,50^.

### scMomer embedding facilitates in-silicon perturbation

Understanding cellular responses to perturbations is critical for studying drug actions, disease mechanisms, and therapeutic interventions. However, the vast combinatorial space of potential perturbations far exceeds the limits of experimental feasibility. With the increasing availability of large-scale perturbation data, computational methods are now widely adopted to simulate gene or drug perturbations and predict gene expression changes under unseen conditions. scMomer offers unique advantages over existing models in this context. First, its transformer-based architecture enables the modeling of long-range gene dependencies, which is important for capturing complex regulatory responses. Second, its dual encoders for RNA and ATAC modalities allow the model to extract distinct yet complementary patterns from each data type. More importantly, as a multi-modal foundation model, scMomer is capable of cross-modality prediction, granting it the unique ability to model perturbation effects across different omics layers, a feature that traditional single-modality models lack.

To evaluate scMomer’s performance in perturbation response prediction, we conducted experiments on two datasets: Adamson^51^ and Norman^52^. Following the setup used in CellFM and scFoundation, we integrated gene embeddings from scMomer into the GEARS^13^ for downstream prediction (Supplementary Fig. 11). We assessed model performance using Pearson correlation on the top 20 differentially expressed genes. Results showed that scMomer outperformed GEARS, achieving improvements of 0.034 in PCC (Supplementary Table 5).

We further explored the cross-modality perturbation prediction capability of scMomer. Using a multi-modal perturbation dataset curated by Wei et al.^53^, we manually held out specific perturbation combinations to simulate in-silico perturbations. Following the process of Luo et al.^28^, we evaluated the performance using averaged pseudo-bulk profiles. The results demonstrated that scMomer could accurately capture changes in chromatin accessibility in response to different perturbation combinations, showing high predictive power for top differential peaks (Supplementary Table 6).

## Discussion

Recent advances in single-cell foundation models are revealing the underlying “language” of cells and molecules. However, existing models built upon single modalities often fall short in capturing the complexity and heterogeneity of biological systems. In this study, we present scMomer, a modality-aware pretraining framework designed to support robust representation learning under missing modality conditions in single-cell multi-omics data. Through a structured three-stage training paradigm, scMomer progressively learns unimodal representations, aligns cross-modal information, and ultimately produces modality-agnostic embeddings via knowledge distillation. This design enables scMomer to generalize effectively to diverse downstream tasks, including gene function prediction, cell type annotation, drug response inference, and perturbation modeling.

Despite its promising performance, scMomer has several limitations. First, the availability of high-quality multi-omics datasets remains limited. Our multi-modal training was conducted using atlas-level pseudo-paired multi-omics dataset, which may bias the learned embeddings toward specific biological contexts and underrepresent rare functional states or disease conditions. Second, scMomer’s performance depends on the architecture choices of its backbone components. Although we demonstrated the feasibility of using scBERT and scCLIP, further improvements could be achieved by exploring more advanced and scalable foundation models.

Several promising pathways deserve future research. The first is to extend scMomer to other types of multi-omics data, such as transcriptome–proteome profiles generated by CITE-seq^54^. The framework may also be adapted to integrate more than two modalities, enabling a more comprehensive understanding of cellular heterogeneity and regulatory mechanisms. Although tri-modal datasets are currently limited, techniques such as optimal transport^55^ and manifold alignment^56^ offer feasible strategies to construct pseudo-matched datasets across modalities. In addition, incorporating Mixture-of-Experts (MoE)^57^ mechanisms to enable flexible input of arbitrary modalities also holds great promise^58^. Another important direction involves enhancing model interpretability to uncover biological mechanisms and identify key molecular features. Gradient-based attribution methods or attention-weight analyses may enable the identification of gene regulatory programs specific to disease states or cellular subpopulations^59^. Overall, as both single-cell profiling technologies and large-scale pretraining methods continue to develop, scMomer provides a generalizable and modular framework for single-cell multi-omics analysis.

## Methods

### Modality-aware pretraining for missing-modality adaptation

A central innovation of scMomer lies in its structured modality-aware pretraining strategy, which enables effective representation learning from incomplete multi-omics data by progressively capturing unimodal dependencies, cross-modal interactions, and modality-agnostic features. The framework consists of three successive stages: modality-specific pretraining, multi-modal fusion, and cross-modality distillation. Each component is designed to reflect biological structure while supporting downstream inference from partially observed inputs.

#### Modality-specific pretraining

We begin with self-supervised learning on each modality to capture internal dependencies. For RNA, we adopt a masked modeling strategy similar to BERT. Specifically, we randomly mask a subset of expressed genes in a cell and task the encoder with reconstructing their expression from the remaining genes. Formally, for a single-cell RNA vector 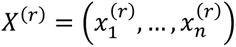, where n is the number of genes, we mask *m*% of the non-zero entries to produce 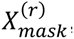, and train the model to predict the original expression values 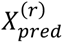. The RNA reconstruction loss is:

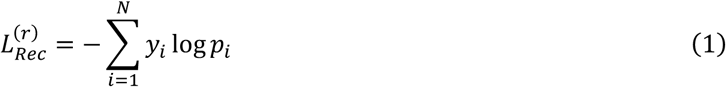

where *N* is the number of masked genes, and *y*_i_, *p*_i_ denote the true and predicted binned expression of gene *i*.

For ATAC, we address the much higher dimensionality (typically over 100,000 peaks) by first segmenting the genome into contiguous patches. Each patch contains a fixed number of peaks mmm, yielding a patch-level representation *X*^(a)^ = (*patch*_1_, …, *patch*_p_). We randomly mask a subset of patches and reconstruct their accessibility using surrounding patches. The reconstruction loss for ATAC is defined as:

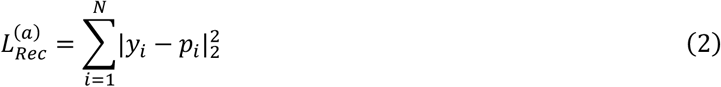

where *y*_i_, *p*_i_ are the true and predicted patch accessibility values.

This stage enables the model to learn intra-modality structures such as gene–gene co-expression and spatial chromatin openness without requiring supervision. Although we adopt masked modeling as the default strategy, the framework remains compatible with alternative self-supervised objectives such as sentence-like gene tokenization or read-depth-aware pretraining.

#### Multi-modal fusion and alignment

In the second phase, we align and integrate paired RNA and ATAC representations to capture cross-modal relationships. Starting from the pretrained modality-specific encoders, we concatenate RNA (*f*^(r)^) and ATAC (*f*^(a)^) embeddings and pass them through a fusion network:

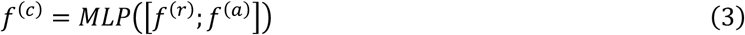

where […; …] denotes vector concatenation.

To ensure biological completeness, we apply reconstruction losses on both RNA and ATAC inputs from the fused embedding *f*^(r)^, forcing the model to retain features relevant to both modalities. Additionally, we introduce a modality matching discriminator that distinguishes whether a given RNA–ATAC pair originates from the same cell. This encourages alignment of paired inputs while preserving biological independence across modalities. The total loss function for this phase is:

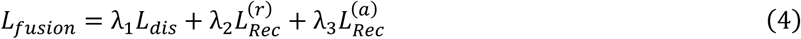

where *L*_dis_ is the binary cross-entropy loss for pair matching; *λ*_1_, *λ*_2_, and *λ*_3_ are hyperparameters to balance each loss.

#### Cross-modality distillation

To support inference under missing modality conditions, we introduce a lightweight cross-modality bridge trained to approximate the missing encoder’s output. Specifically, we simulate scenarios where only RNA is available, freeze the RNA encoder, and train a student model to predict the ATAC representation learned in the previous phase. Let 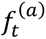 denote the ATAC embedding from the pretrained encoder, and 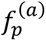 the predicted embedding from the bridge network. The distillation loss is:

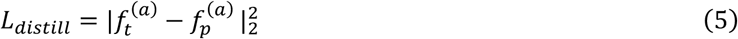

We additionally reconstruct the RNA and ATAC profiles from the fused embedding to preserve biological meaning. The total loss is:

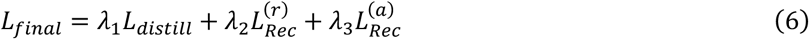

This distillation process equips scMomer with the ability to approximate multi-modal cell embeddings using only unimodal inputs, enabling robust analysis in real-world datasets where only RNA or ATAC profiles are available.

### Overview of scMomer architecture

scMomer adopts a modular and flexible architecture designed for single-cell multi-omics representation learning. Its components include modality-specific encoders for RNA and ATAC data, modality-specific decoders, a cross-modal integration module, and a lightweight feedforward-based bridge for cross-modality translation. This design ensures that each module can be independently improved or replaced, allowing future integration of more advanced backbones or modalities.

#### RNA encoder

The RNA encoder is based on scBERT, a pre-trained transformer model tailored for modeling gene expression. It consists of a gene tokenizer, an expression tokenizer, and a multi-layer Performer encoder^60^. Gene identities are first mapped to vector representations using gene2vec embeddings, which are trained to capture gene co-expression structure. Expression values are then discretized into bins, reflecting relative expression levels, and embedded separately. The final input embeddings are the sum of gene and expression embeddings. To efficiently handle the long input sequences typical of single-cell transcriptomes, we employ the Performer architecture, which approximates full self-attention with linear complexity. This enables modeling of long-range dependencies across thousands of genes per cell. After passing through multiple layers of Performer blocks, the output is projected via a 1D convolution and a linear layer to yield the RNA-based cell embedding.

#### ATAC encoder

For ATAC encoder, we use a ViT-based encoder^31^ adapted from scCLIP to account for the high dimensionality of chromatin accessibility data. The genome is segmented into ordered, fixed-length “patches” based on genomic position, analogous to image patches in vision transformers. Each patch represents a local region of chromatin accessibility and is embedded accordingly. We also add 1D positional encodings to preserve spatial relationships along the genome. These patch embeddings are then passed through a Performer encoder and pooled into a fixed-length ATAC-based cell embedding.

#### Integration module

To form joint multi-omics representations, the RNA and ATAC embeddings are concatenated and passed through a linear projection. This projection forms the integrated cell embedding, which captures complementary information from both omics layers. The integration module is lightweight and modality-agnostic, enabling efficient downstream usage and flexible extension to additional modalities.

#### Decoders and cross-modality bridge

scMomer includes two modality-specific decoders, each implemented as a simple feedforward neural network (FFN). These decoders are used for reconstructing masked features during pretraining and are intentionally kept lightweight to isolate the quality of the learned embeddings. To support inference under missing modality conditions, scMomer incorporates a cross-modality bridge module, also built using FFNs. This student model is trained to recover missing modality embeddings (e.g., ATAC) from the available RNA-based embedding. The simplicity of the FFN structure, as shown in prior work such as scPair, is sufficient for effective translation when the input representation is biologically meaningful.

### Datasets

scMomer is pretrained in a self-supervised manner on large-scale single-cell multi-omics data and evaluated across a range of downstream tasks under missing modality settings. To avoid data leakage, we strictly separate training and evaluation datasets. During pretraining, we utilize high-coverage atlas-scale datasets to enhance model generalization. For downstream evaluation, we adopt high-confidence, task-specific datasets to ensure rigorous assessment.

#### Multi-modal datasets

We use two single-cell multi-omics datasets for model pretraining and cross-modality evaluation: a fetal multi-organ atlas and a human brain dataset. Due to the current lack of large-scale real paired multi-omics data, we adopt a curated pseudo-paired dataset from Xiong et al.^22^, in which cells are manually matched based on annotated cell types. The fetal atlas dataset contains 377,134 cells across 47 cell types, with 36,601 genes and 1,154,464 ATAC peaks. The human brain dataset consists of 3,233 healthy brain cells spanning 7 major cell types, each with paired gene expression and chromatin accessibility profiles.

#### Disease datasets

To evaluate the biological relevance and integrative capacity of scMomer’s representations, we adopt two disease-focused single-cell datasets curated by Chen et al.^34^ The aorta dataset^61^ includes 10 cell types from 11 patients with distinct pathological conditions, including healthy, ascending thoracic aortic aneurysm (ATAA), ATAA with root aneurysm, and ATAA with descending thoracic involvement. The cardiomyocyte dataset^62^ contains three cardiac disease subtypes: non-failing (NF) hearts, hypertrophic cardiomyopathy (HCM), and dilated cardiomyopathy (DCM). Both datasets are subsampled versions of larger cohorts to ensure consistency and computational efficiency.

#### Cell type annotation datasets

For evaluating model robustness in cell type annotation tasks, we used both intra-dataset and out-of-distribution settings. The intra-dataset setting included Zheng68K and multiple pancreas datasets, which are widely used benchmarks. Zheng68K contains 68,450 peripheral blood mononuclear cells (PBMCs) spanning 11 cell types, and is considered challenging due to its high class imbalance and closely related cell populations. The pancreas benchmark consists of four datasets (Baron, Muraro, Segerstolpe, and Xin) collected using different sequencing platforms. To create an out-of-distribution evaluation, we extracted four shared cell types (alpha, beta, delta, and gamma cells) across pancreas datasets. The Baron dataset served as the training set, while the others were used for testing.

#### Gene datasets

To test scMomer’s ability to capture gene-level properties, we used datasets spanning various biological tasks. Gene function prediction tasks were derived from Theodoris et al.^15^, involving binary classifications such as dosage-sensitive versus insensitive transcription factors, bivalent versus non-methylated genes, and long-range versus short-range regulators. For gene-gene interaction (GGI) prediction, we used a large-scale dataset from Du et al.^35^, containing over 200,000 annotated gene pairs with binary labels. Protein-protein interaction (PPI) prediction was performed using the HuRI dataset curated by Luck et al.^37^, which similarly includes binary-labeled interactions between protein pairs.

#### Drug response datasets

To evaluate the applicability of scMomer in pharmacogenomic tasks, we tested its performance on both single-drug response prediction and drug combination modeling. For single-drug response prediction, we adopted a dataset curated by Liu et al., which integrates pharmacological and genomic profiles from three publicly available resources: GDSC, CCLE, and TCGA. This dataset includes gene expression profiles of 561 cancer cell lines covering 31 cancer types, along with response data for 223 compounds. Molecular features of the drugs were also incorporated to facilitate learning drug–cell line interactions. For drug combination prediction, we used the dataset from O’Neil et al., which includes annotated drug pairs and their combinatorial effects. SMILES representations of compounds were retrieved from PubChem based on drug names, and corresponding gene expression profiles were extracted from the CCLE database.

#### Perturbation datasets

To assess scMomer’s ability to capture regulatory changes under genetic perturbations, we employ two large-scale Perturb-seq datasets. The Adamson dataset, comprising 87 single-gene perturbations, each with ∼100 perturbed cells and >7,000 control cells. The Norman dataset, containing 105 single-gene and 131 combinatorial perturbations, enabling analysis of synergistic and antagonistic effects in gene regulation.

### Data pre-processing

We followed the data processing pipeline of scBERT. For a gene expression matrix, log transformation was first performed using a size factor of 10,000. Subsequently, quality control was carried out by retaining only cells with at least 200 expressed genes.

### Downstream applications

To comprehensively evaluate the advantages of scMomer in downstream tasks, we conducted experiments across both gene-level and cell-level prediction tasks. Unlike unimodal models, scMomer leverages multi-modal priors to learn representations that encode both within-modality structures and cross-modal relationships. We compared scMomer against a series of baseline methods, including both general-purpose single-cell embedding models and task-specific predictors. All baseline models were trained using their default configurations. For models such as scBERT that do not natively provide cell-level embeddings, we followed the strategy in Hao et al.^16^ and aggregated gene embeddings using both max and average pooling to construct cell-level representations.

#### Gene-level tasks

To construct a unified gene embedding space, we computed gene-level representations by averaging the embeddings across all cells in which each gene was observed. We selected the intersection of gene sets shared across all models to build a consistent evaluation dataset. For functional gene property prediction, we adopted four binary classification tasks as described in Theodoris et al.^15^ These tasks were evaluated using logistic regression and random forest classifiers implemented in scikit-learn, with 5-fold cross-validation and a train-test split ratio of 8:2. Let *e*_i_ ∈ ℝ^d^ denote the embedding of gene *i*. For gene–gene interaction (GGI) prediction, we constructed input features by concatenating embeddings of gene pairs, i.e., *Z*_ij_ = [*e*_i_; *e*_j_] and trained a binary classifier to predict interaction labels. For protein–protein interaction (PPI) prediction, we first mapped protein identifiers to gene symbols using UniProt^36^, then followed the same procedure as in the GGI task.

#### Cell type annotation

To evaluate cell-type classification performance, we used a three-layer feedforward neural network as a classification head. Given a cell embedding *h*_c_, the model predicts cell type probabilities via 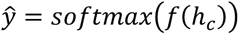, where *f*(⋅) denotes the feedforward layers. Cross-entropy loss was used to optimize the model during training.

#### Drug response prediction

For drug response prediction, we followed the experimental setup from DeepCDR^46^. Specifically, we randomly held out 5% of the dataset for testing. The original DeepCDR model uses gene expression profiles as input features, excluding mutation and methylation information. To enable a fair comparison, we replaced the gene expression profiles with cell line embeddings derived from scMomer while keeping all other model configurations unchanged. For drug synergy prediction, we adopted the DeepSynergy^63^ framework. Drug–cell pair features were constructed by concatenating the drug representations and the gene expression-based cell embeddings.

#### Perturbation prediction

We evaluated perturbation response prediction following recent benchmarks in large-scale single-cell foundation models, such as scFoundation^16^ and CellFM^64^. Specifically, we used the GEARS^65^ as the backbone and replaced its native gene embeddings with those derived from scMomer. The model was trained to predict gene expression shifts under perturbation conditions.

## Data availability

All data used in this study are publicly available. Fetal atlas dataset can be accessed at https://github.com/jsxlei/scCLIP. Human brain dataset is available at https://cf.10xgenomics.com/samples/cell-arc/2.0.0/human_brain_3k/human_brain_3k_filtered_feature_bc_matrix.h5. Zheng68K dataset is available at https://github.com/TencentAILabHealthcare/scBERT.

Pancreas data is available at https://hemberg-lab.github.io/scRNA.seq.datasets/. Gene-level dataset is available at https://github.com/yiqunchen/GenePT. Drug response dataset can be found at the DeepCDR repository: https://github.com/kimmo1019/DeepCDR. Perturbation dataset is available from the GEARS repository: https://github.com/snap-stanford/GEARS. Cross-modality perturbation data from PerturBase^53^ is available at http://www.perturbase.cn/browser/19.

## Acknowledgment

This work is supported by the National Natural Science Foundation of China (No. 62322112), and the Science and Technology Development Fund of Macau (No. 0133/2024/RIB2).

## Competing interests

The authors declare no competing interests.

## Author contributions

L.W., R.S., and Q.Z. conceived the study. Y.L. led the analysis. All of the authors contributed to writing the paper and have approved the manuscript.

## Notes

### Competing Interest Statement

The authors have declared no competing interest.

